# Ongoing evolution of KRAB zinc finger protein-coding genes in modern humans

**DOI:** 10.1101/2020.09.01.277178

**Authors:** Christian W. Thorball, Evarist Planet, Jonas de Tribolet-Hardy, Alexandre Coudray, Jacques Fellay, Priscilla Turelli, Didier Trono

## Abstract

**Background:** Krüppel-associated box (KRAB) zinc finger proteins (KZFPs) constitute the largest and fastest evolving family of gene regulators encoded by the human genome. Recent data indicate that many KZFPs serve as repressors of transposable element-embedded regulatory sequences (TEeRS) and that the evolutionary turnover of KZFP genes is mainly attributable to the changing transposable element (TE) load of their hosts. However, how natural selection and genetic variation are shaping this process is still poorly defined.

**Methods:** Genetic information was collected from nine primate species and 138,500 human genomes. Gene-wide as well as functional amino acid position specific constraint was calculated across all human KZFPs.

**Results:** We found that the most conserved KZFPs, some of which go back close to 400 million years, have been subjected to marked negative selection in the evolutionarily recent past and are very homogeneous within the human population. In contrast, younger, largely primate-restricted family members present evidence of less negative selection than the rest of genome and lower levels of coding constraint, particularly within the sequences encoding the functional sites of their zinc finger (ZF) arrays. We defined 33 sets of KZFP paralogs, which pairwise displayed a broad range of coding constraints differentials, with more recently emerged paralogs usually displaying a higher frequency of putatively deleterious mutations and missense variants within the functional sites of their ZF arrays than their source gene. Finally, we identified three KZFP genes more constrained in the genomes of individuals of African ancestry than in Europeans, with their modes of expression or DNA targets pointing to possible links between these inter-populational genetic differences and regional differences in the prevalence of some diseases.

**Conclusions:** This work shows how the ongoing selection of KZFPs contributes to modern human genetic variation, in particular through the constraint of putatively deleterious- and missense variants in functional protein sites, and how ongoing interplays between environment and KZFP genes might be impacting the biology of modern humans.

## Background

KRAB-containing zinc finger protein (KZFP) genes emerged approximately 420 million years ago (MYA) in the last common ancestor of coelacanth (*Latimeria chalumnae*), lungfish and tetrapods, and have expanded since then by gene and segment duplication to count more than 350 representatives in the human genome [1,2]. Their products are characterized by an N-terminal KRAB domain and a C-terminal array of two to more than forty Cys2-His2 (C2H2) zinc fingers (ZFs) with sequence-specific DNA binding potential [3]. Close to 45% and 90% of human KZFPs are primate- and mammalian-restricted, respectively. Most of these evolutionarily more recent KZFPs act as repressors of the transcriptional activity of transposable elements (TE)-embedded regulatory sequences (TEeRS) via the KRAB-mediated recruitment of KAP1 (KRAB-associated protein 1, also known as TRIM28), which serves as a scaffold for a heterochromatin-inducing complex [1,4]. The primary consequence of this effect is not to prevent further transposition since mutations inactivate most TEs targeted by KZFPs, but to facilitate the cooption of TEeRS into transcriptional networks [5]. In contrast, more conserved KZFP family members, which are often endowed with variant KRAB domains, do not recruit KAP1 but display other protein interactomes suggestive of functions such as maintenance of genome architecture or RNA processing [4]. Individual members of the KZFP family have been implicated in processes as diverse as adipogenesis, cell differentiation, genomic imprinting, metabolic control or meiotic recombination [6—14]. Since KZFPs from coelacanth, our most distant KZFP-encoding relative, bind the cognate KAP1, it is likely that KZFPs first emerged as TE-controlling repressors and from then on underwent continuous selection through turnover of their hosts’ TE loads. While derivatives were occasionally produced that escaped this evolutionary flushing by development and exaptation of novel functions, the largest fraction of the KZFP gene pool of individual lineages is renewed over time, contributing to the species-specificity of regulatory networks. How much genetic drift occurs on a much shorter time scale, for instance since modern humans emerged and spread over the globe, is nevertheless unknown, which prompted the present study.

The recent availability of large human genome sequencing datasets such as the Genome Aggregation Database (gnomAD) [15] provides an opportunity to identify biologically essential human genes through their depletion in protein-altering variants, also known as coding constraint [16–20]. However, while gene-wide measures of constraint are useful for identifying haploinsufficient genes and for interpreting disease-associated variants, they do not capture biologically important, site-specific constraints within protein domains. The DNA-binding specificity of KZFPs is dictated mainly by three amino acids in the alpha-helical region of each ZF, known as the zinc fingerprint [1]. Thus, missense variants in these positions alter the target preference and thereby the functional impact of a KZFP. At the same time, changes in the two cysteine or two histidine residues (C2H2) that stabilize each ZF through the recruitment of a zinc ion inactivate these structural domains altogether. By combining phylogenetics and large-scale population genetics, the present work examines recent changes of the KZFP gene pool in the human lineage. Its results strongly suggest that a large fraction of KZFP genes are subjected to ongoing selective pressures, and in return contribute to phenotypic differences between individuals.

## Methods

### Map of the human KRAB Zinc Finger protein clusters

KZFP pairs were detected and their age defined as described in [1]. In short, the human genome (hg19) was translated in 6 reading frames and scanned for zinc finger and KRAB domains using Hidden-Markov-Models (Pfam [21]: KRAB (PF01352) and zf-C2H2 (PF00096)). Hits for KRAB and Zinc Finger domains were combined based on proximity and strandness and then manually curated and integrated with existing gene or pseudogene annotations. Their age is based on sequencing similarity with orthologues in other species. For this analysis the position of each KZFP was reduced to a point located in the center between the KRAB and Zinc Finger domains. Alternative chromosomes were ignored. The KZFP clusters were defined as having at least 3 KZFPs that are no more than 250 kb apart from the center of another member, consistent with [22]. The clusters are named after their chromosome and then numbered starting from the short arm of the chromosome. The size of chromosomes and positions of centromeres were taken from UCSC genome browser annotation data [23] for hg19.

### Primate phylogeny and natural selection

The time of divergence (i.e. branch lengths) between human, chimpanzee, gorilla, orangutan, macaque, marmoset, tarsier, galago (a.k.a. bush baby) and mouse lemur was obtained from 10KTrees, which uses Bayesian inference to estimate these [24].

Measures of natural selection in terms of dN/dS across the nine primate species listed above was obtained with PAML (v4.4) as previously described [25].

### Human genetic variation data

Human genetic exome and whole genome sequencing data were obtained from The Genome Aggregation Database (gnomAD) [16,26] (release-2.0.2) for 123,136 and 15,496 individuals, respectively. The released genetic data was processed and filtered through several steps to guarantee that only high-quality variants were included. First, all variants +/- 1kb around the KZFP canonical transcripts as defined by Ensembl (v75, hg19) were extracted and filtered for variant quality, thus only retaining variants annotated as “PASS”. Second, all indels were normalized and multiallelic variants split using BCFTOOLS (v1.8) and reannotated with the Variant Effect Predictor [27] and LOFTEE (v0.3beta). Third, all missense and loss of function (LoF) variants, defined as either frameshift, stop-gain or splice variants, were extracted from both the exome and whole genome datasets and either low confidence or flagged LoF variants were removed. The latter was primarily due to LoF variants found in the last 5% of the canonical transcript. Since genomic sequencing methods can yield variable coverage of genetic regions, especially when it comes to exome sequencing that is dependent on the capture of previously annotated protein-coding genes, we excluded all canonical transcripts having an average per-base coverage < 20x. Thus, bringing the total number of included KZFPs to 361. Furthermore, exons with an average per-base coverage < 20x were also removed, and the lengths of the coding sequences used later for normalizations were adjusted accordingly. Finally, the filtered exome and genome datasets were combined, and the allele counts and frequencies for all variants were recalculated, prior to the removal of all singletons (allele count = 1) to hinder inflation of observed mutational events due to potential technical artifacts.

### Domain and site specifications

The genomic positions of the C2H2 zinc finger domains were obtained from the Ensembl database (v75, hg19). For each KZFP, only the ones from the canonical transcripts (as defined by Ensembl) were considered. The positions of the specific amino acids within these domains were computationally annotated.

Z scores for the cysteine and histidine (C2H2) residues and the DNA fingerprint positions within the ZF domains were calculated with the number of missense variants normalized to the number of ZF domains within the canonical transcript of each KZFP. For missense and LoF variants spanning either the whole CDS or a full protein domain, the number of variants per gene, x, was normalized by the length of the canonical coding sequence prior to Z score transformations.

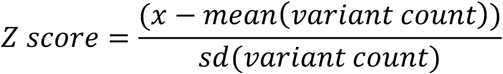

### Zinc fingerprint evolutionary conservation score

To distinguish the evolutionary conservation of each KZFP within paralog pairs and determine the source gene of the pair, each array of zinc finger DNA-binding amino acids were concatenated as a single sequence, separated by an artificially added character. To align, we used a customized version of Blosum80 matrix (https://www.ncbi.nlm.nih.gov/IEB/ToolBox/C_DOC/lxr/source/data/BLOSUM80), modified by adding a +100 score when the special character was matched by itself, and a −100 penalty when the special character was matched by any amino acid (but not by a gap). Pairs of Zinc Finger Arrays were then aligned globally with pairwise2.align from Biopython [28] using the modified matrix, an opening gap penalty of −20, a gap extension penalty of 0 and no penalty for end gaps. For each alignment, the score was normalized by the alignment length and corrected for the participation of the special character. To compute the final score, we then decided to not use only ratio of aligned AA in the sequence (amino acid level), but also to incorporate the ratio of fully aligned zinc fingers (zinc finger level), with weights of 0.5 to each terms.

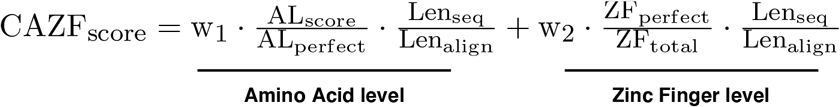

With ALscore being the Blosum80 alignment score of the alignment, ALperfect the score of the alignment of the sequence against itself, Lenseq being the length of the sequence, Lenalign the length of the alignment (= the longest of both sequence), ZFperfect the number of zinc finger that aligned without mismatches and ZFtotal the total number of Zinc Finger contained in a given KZFP.

### ChIP-exo data

ChIP-exo data in 293T cells was obtained from [1] using overexpression of tagged KZFPs. Reads were mapped to the human genome assembly hg19 using Bowtie2 short read aligner [29], using the --very-sensitive-local. Peaks were called using MACS v1.4.2.1 [30] with defaults parameters with control as in [1] and with the parameter --keep-dup all. For the rest of the analyses, only peaks with MACS score > 80 were kept.

KZFP motifs were identified using RSAT and the peak-motifs function [31].

TE enrichment analyses from ChIP-exo data was performed using TEnrich (https://github.com/alexdray86/TEnrich). TEnrich relies on a binomial exact test, which compares the number of peaks on a TE family to what would be expected by random while taking the total genomic span of each TE family into account. A peak is considered to be on a TE when the overlap is above 50%.

Gene ontology (GO) enrichment analyses from ChIP-exo data of ZNF114 and ZNF714 for biological processes was performed using GREAT (v4.0.4) [32], using the default settings and significance levels.

### Gene expression data and correlations

Median gene expression counts across 40 tissues were obtained from GTEx (v7) [33]. The median expression value of the various brain sections included in GTEx were combined into a single category named “Brain”. KZFPs were considered expressed when transcripts per million (TPM) were above 0.3 TPM. Spearman correlations were used to assess the expression similarities of KZFP paralogs.

### Statistics

All statistical analyses were performed using R (v3.6.0). Wilcoxon rank sum tests were used for all paired analyses (e.g, primate vs. non-primate groups) and correlations were made using Spearman’s rank correlation coefficient.

## Results

### Genomic distribution and recent evolution of human KZFP genes

The KZFP gene family has been expanding primarily through gene or segment duplications, with minor contributions of translocation, retrotransposition, or recombination. As a consequence, KZFP genes are often grouped in clusters, with adjacent family members encoding proteins with significant levels of amino acid similarity. We established a census of the chromosomal distribution of 466 juxtaposed KRAB and C2H2 poly-zinc finger domains identified in the human genome, independent of their presence or not in an annotated gene (Figure 1). We found 74% (n = 344) of these domain pairs to be located within one of 30 clusters, defined here as a group of at least three KRAB/poly-ZF couples separated by less than 250 kb. The majority (n = 248) concentrated in 11 clusters on chromosome 19. Using previously estimated evolutionary ages [1], we further determined that isolated protein-coding KZFP genes tended to be older than their cluster-associated counterparts (p = 3.5e-6). This fits with the proposal that chromosome 19 is the main region of emergence of new KZFP genes [34] and suggests that escaping the tumultuous environment of this chromosome facilitated the fixation of older family members. An exception is the grouping of some of the most ancient KZFPs, namely *ZNF282, ZNF777*, and *ZNF783*, in cluster 7.3 at the distal end of human chromosome 7 long arm as previously noticed [35].

**Figure 1.**
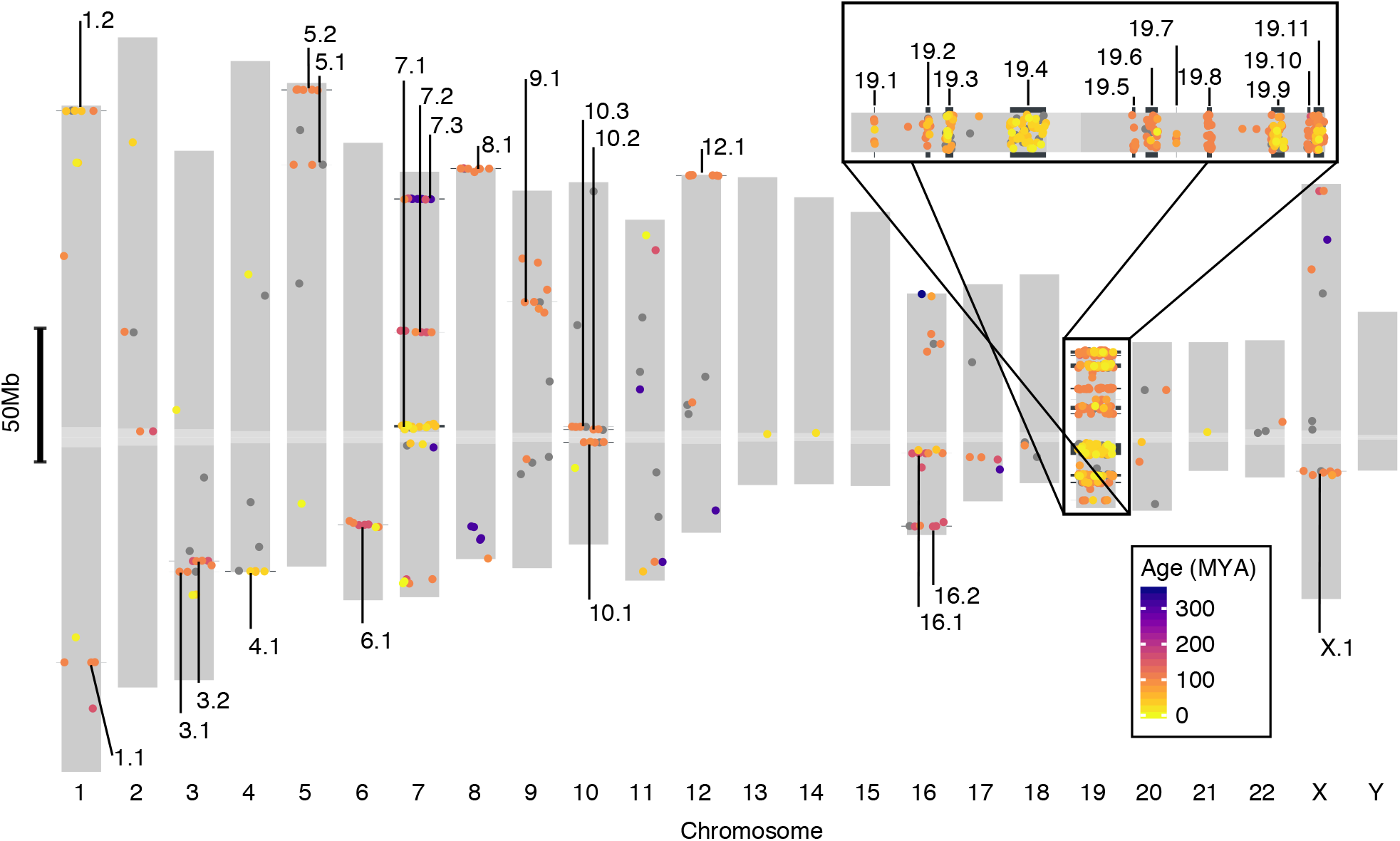
Genomic distribution of KZFP genes. Dots indicate relative chromosomal position of KZFP genes (defined by juxtaposed KRAB- and ZF-coding domains), with color code indicative of age (grey for unassigned) and numbered clusters pointed to in black. Higher magnification of chromosome 19 is presented on top right.

To complement this first analysis, we examined the recent evolution of KZFP genes in the primate lineage as described previously [25]. For this, we determined their gene-wide ratio of non-synonymous (missense) (dN) to synonymous (dS) substitutions (dN/dS), based on genome sequence data from human, chimpanzee, gorilla, orangutan, macaque, marmoset, tarsier, galago (a.k.a. bush baby) and mouse lemur, that is, over ~6 to ~74 million years of divergence (Figure 2A). We found KZFPs to have significantly higher dN/dS values than genes coding for other proteins (p = 1.46e-49), including KRAB-less ZF proteins (p = 3.04e-46) (Figure 2B), confirming previous observations [35,36]. We also noted that the distribution of dN/dS values was bimodal amongst KZFPs, with younger, primate-specific genes displaying higher scores than family members having emerged earlier in evolution (p = 2.69e-21) (Figure 2C) and the dN/dS ratio of KZFP genes being anti-correlated with their estimated age (rho = −0.61) (Figure 2D).

**Figure 2.**
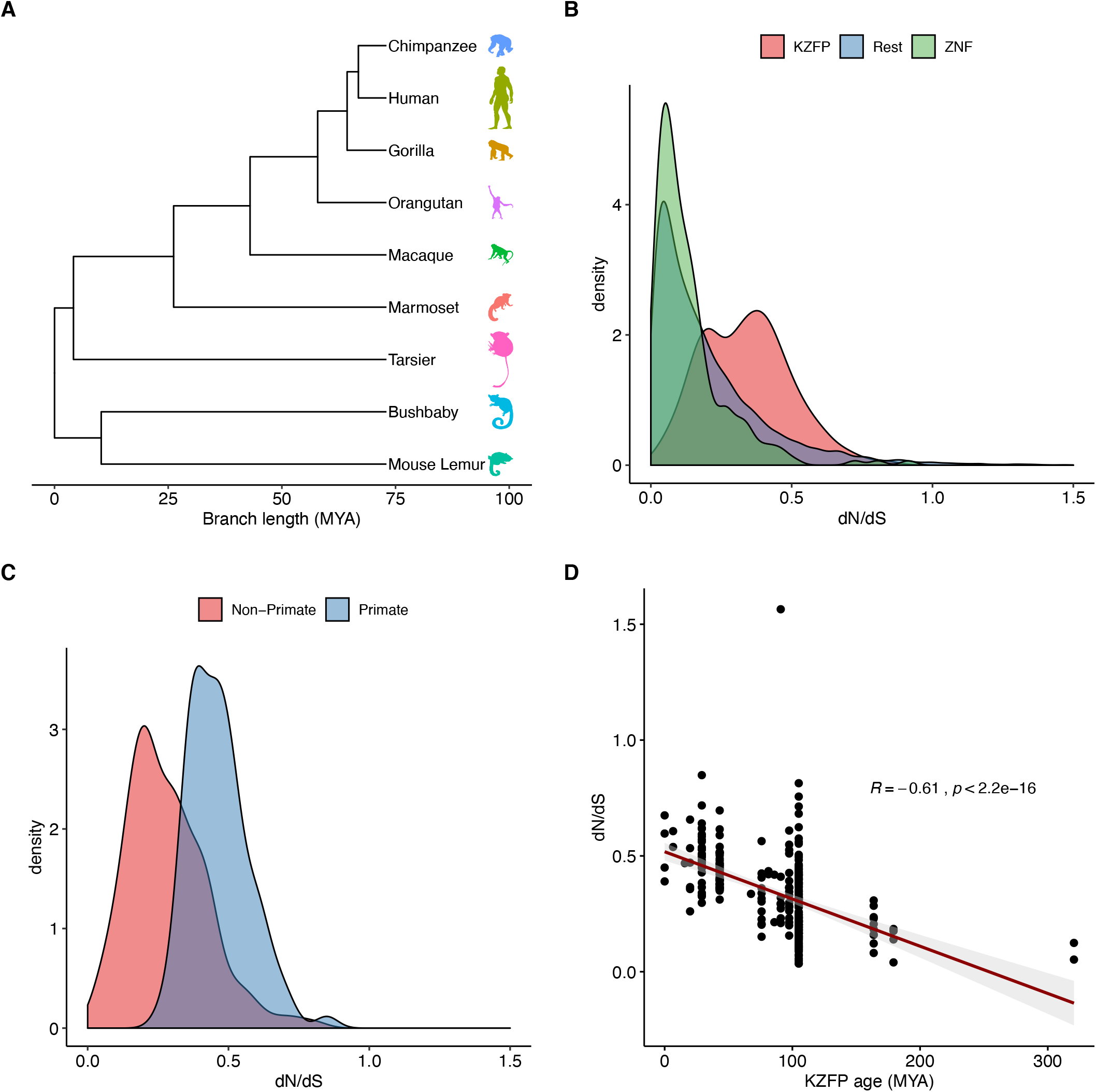
Evolution of human KZFP genes in primate lineage. (A) Phylogenetic tree of primate species used to calculate natural selection of human genes, with branch length indicating approximate time of divergence in million years (MYA). (B) Distribution of PAML dN/dS values of natural selection for KZFPs (red), non-KRAB ZFPs (green) and all remaining genes in the genome (blue). (C) dN/dS distribution of primate-specific KZFPs (blue) and older (red) KZFP genes. (D) Spearman correlation of the dN/dS values and estimated age of KZFP genes.

### Conservation-related gradient of human KZFP polymorphisms

The coding constraint of a gene or fragment thereof reflects the strength of selective pressures imposed on its sequence, hence the relative functional importance of the corresponding protein or protein domain for a given species. Typically, highly constrained coding regions correspond to loci where mutations are either associated with disease or are completely absent because they cause sterility or embryonic lethality. To examine the coding constraints imposed on human KZFP genes, we examined genetic variation amongst 138,632 individuals (15,496 genomes and 123,136 exomes) cataloged in gnomAD v.2.0.2. After removing coding sequences with low coverage and dismissing singletons to reduce the impact of false positives resulting from sequencing or alignment errors, we extracted protein-altering variants (missense and predicted loss of function -LoF-by frameshift, gain of stop codon or alteration of essential splice sites) within the canonical transcripts of all remaining KZFPs (n = 361). For the estimation of gene-wide constraint, we normalized the number of variants for the length of the canonical coding sequence and translated the result into a z score to standardize values (Figure 3AB). Accordingly, negative deviation from the mean was a sign of increased purifying selection as a consequence of reduced frequency of protein-altering variants. However, we did not correct for a theoretically expected number of mutations as frequently done in this type of analysis because the unstable structure of the ZF array-coding region of KZFP genes renders this parameter unpredictable [2]. Gene-wide, LoF and ZF domain-specific scores modestly correlated with previously measured dN/dS ratios and with the age of the KZFPs (Figure 3C). Examining individual domains revealed that this association stemmed mainly from the ZF C2H2- and to a lesser extent, fingerprint-coding sequences. Of note, other codons of the ZF-coding regions displayed no significant constraint, confirming that essential positions in ZFs are limited to the structure-conferring cysteine and histidine residues and the target-defining fingerprint residues at positions −1, +3 and +6 of the ZF alpha helix [37].

**Figure 3.**
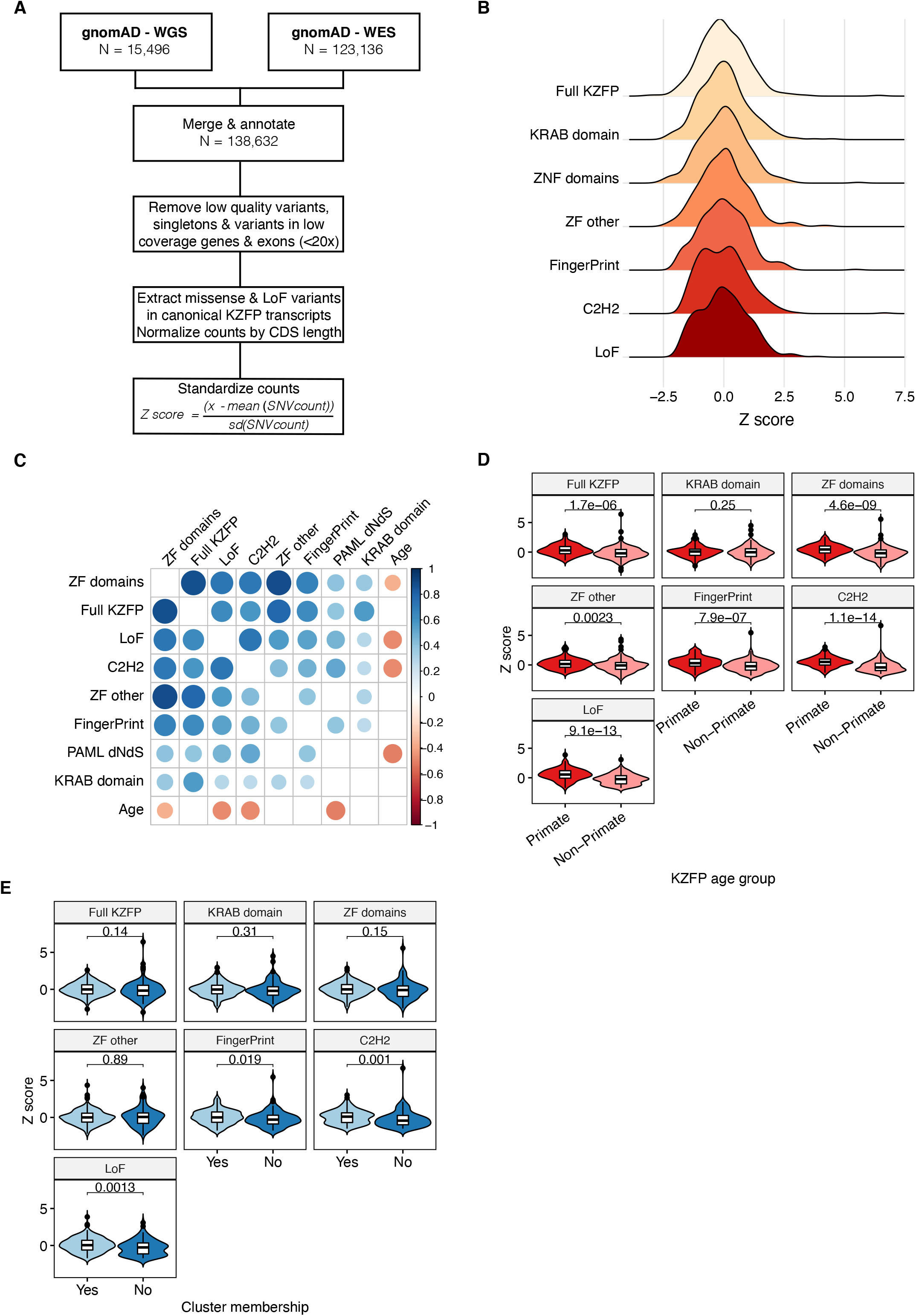
Coding constraints of KZFP genes. (A) Schematic of genetic constraint Z score calculation. WGS/WES, whole genome/exome sequencing; LoF, loss of function variant; CDS, coding sequence. (B) Distribution of indicated Z scores; lower score indicates increased constraint compared to average of all KZFPs. Full KZFP, all variants within the canonical KZFP transcript; KRAB domain, only variants in the KRAB domain; ZF domains, variants within the ZF domains; ZF other, variants in non-functional positions within the ZF domains; FingerPrint, variants in the ZF fingerprint positions; C2H2, variants in the cysteine or histidine positions of the ZF domains; LoF, loss of function variants (C) Spearman correlations between Z scores, level of natural selection (PAML dN/dS) and estimated age of KZFPs. The colors and their intensity represent the direction and strength of the correlations, with blue representing a positive- and read a negative correlation. Only significant correlations after Bonferroni correction are shown. (D) Primate vs. non-primate KZFP constraint across indicated KZFP domains or residues. (E) Relative constraint of indicated regions for KZFPs inside vs. outside clusters.

Primate-restricted, younger KZFPs were significantly less constrained both in terms of LoF (p = 9.1e-13) and missense variation (p = 1.7e-6) than their older counterparts (Figure 3D), with the difference mainly residing in sequences coding the poly-ZF (p = 4.6e-9) rather than the KRAB domain (p = 0.25). Within ZFs, the C2H2- and fingerprint-defining positions were again the most influential (p_ZFc2h2_ = 1.1e-14 and p_ZFprint_ = 7.9e-7), compared to the other non-functional positions of the ZF domains (p_ZF other_ = 0.002). Correlating with their age, isolated KZFP genes were more constrained at sequences encoding the ZF C2H2 residues (p = 0.001) and fingerprint-defining positions (p = 0.02) and displayed lower LoF scores (p = 0.001) than their cluster-associated counterparts, consistent with their stabilization over longer evolutionary times (Figure 3E). However, coding constraints were also highly heterogeneous within most clusters, indicating that differential selective pressures are rapidly exerted on members of a same cluster (Figure S1).

No LoF variants were detected amongst all examined individuals for *ZFP92, ZNF606, ZNF81, ZNF777, ZNF121, ZNF250*, and *ZNF597*. The *ZFP92, ZNF81*, and *ZNF777* genes were also devoid of any missense mutations in their C2H2- or fingerprint-coding positions, while some were detected in *ZNF121, ZNF250, ZNF597*, and *ZNF606* albeit at extremely low allele frequencies. For a majority of other KZFPs (n = 213), some heterozygous but no homozygous LoF variants were observed. Nevertheless, a significant number (n = 148) presented homozygous LoF variants in at least two individuals, suggesting reduced constraint (p < 2.22e-16) (Figure S2). On average, members of this subgroup had a younger estimated age than the rest of the KZFPs (p = 2.9e-13).

### Differential coding constraints of human KZFP paralogs

We examined coding constraints amongst 33 identified sets of KZFP paralogs, which we defined as products of gene duplication with ≥ 60% similarity between their zinc fingerprints [1]. Accordingly, for 28 of the 33 paralog pairs, all paralog pair members were located within the same chromosomal cluster. Significant differences were noted, especially at the C2H2-coding positions, with some pairs of paralogs displaying closely similar coding constraints (e.g., *ZNF75A* and *ZNF75D*) while others were markedly divergent (e.g., *ZNF160* and *ZNF665*) (Figure 4A and Figure S3). The level of divergence was not related to the age of the paralog pairs, whether at C2H2-coding- (p = 0.21) or fingerprint-defining positions (p = 0.27). However, the more constrained paralog within a pair was usually also the most conserved in evolution (Figure 4A). For instance, the ~90-million-year old (MYO) *ZNF160* was markedly more constrained than its ~29 MYO *ZNF665* paralog (Figure S4), both at C2H2-coding positions (Figure 4A) and across other features (Figure 4B). A closer examination of *ZNF160* and *ZNF665* zinc fingerprints revealed that some ZFs were completely constrained in both KZFPs, while others were more flexible (Figure 4C). ChIP-seq analyses confirmed that these proteins recognized closely related sequence motifs (Figure 4D) in overlapping sets of genomic targets (Figure 4E), notably some LINE1 integrants (Figure S5). Furthermore, the two paralogs were noted to have roughly similar expression patterns of across 40 tissues according to the GTEx database (rho = 0.89). More globally, we observed that more constrained KZFPs were generally expressed at higher levels and more ubiquitously than their more flexible counterparts (Figure S6), in line with previous reports [16,38].

**Figure 4.**
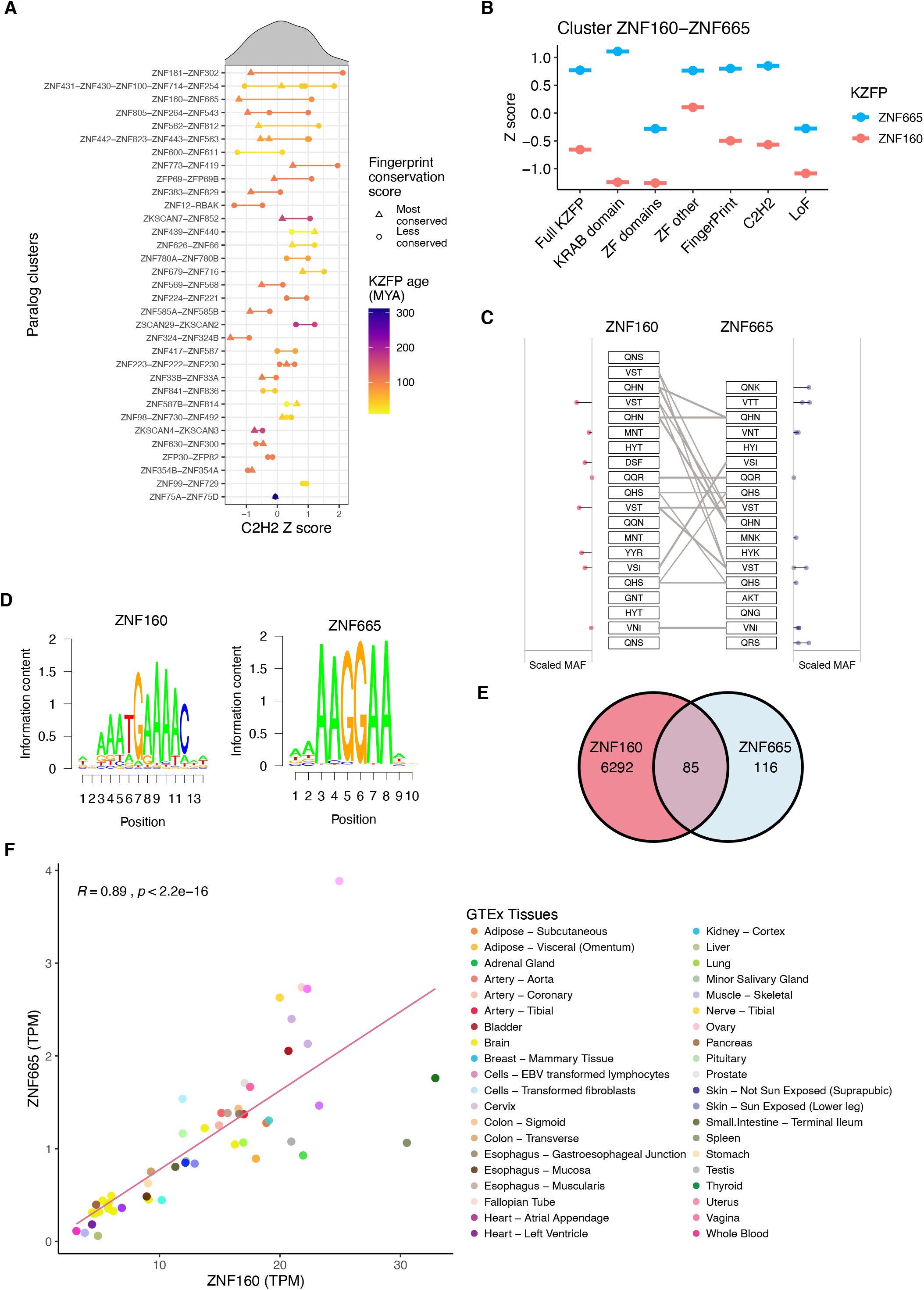
Analysis of KZFP paralogs. (A) Distribution of C2H2 constraint Z scores for indicated sets of KZFP paralogs, arranged from top to bottom according to difference within pairs. Each KZFP is colored according to their respective age, with the line separating them colored as the mean age of the pair. The paralog within each pair with the most conserved fingerprint across evolutionary time, as determined by their CAZF value, is marked by a triangle, while less conserved KZFPs are marked by a dot. The order of the y-axis labels corresponds to the order of the colored points on the graph. (B) Differential constraint Z scores for indicated domains of paralogs *ZNF160* and *ZNF665*. (C) Zinc fingerprints of *ZNF160* and *ZNF665* with the scaled minor allele frequency (MAF) of identified missense variants indicated on the sides. Grey lines indicate identical zinc fingerprints (D) Consensus DNA binding motifs of ZNF160 and ZNF665. (E) Venn diagram of ChIP-exo peaks of ZNF160 and ZNF665 in 293T cells. (F) Expression levels in transcripts per million (TPM) of *ZNF160* and *ZNF665* across all tissues depicted in GTEx.

### Evidence of modern selective pressures on human KZFP genes

To test whether KZFP constraints vary across human populations due to recent selective pressure, we extracted the variant information from the two largest subsets of individuals in the gnomAD database, i.e., 63,369 non-Finnish Europeans (NFE) and 12,020 African/African-Americans (AFR). We calculated their population-specific constraint, adjusting for the difference in sample size to reduce the bias in variant discovery power. Correlation of missense constraints across all KZFP genes was strong (rho = 0.70, Figure S7A), but it was weaker at C2H2- (rho = 0.52, Figure S7B) and zinc fingerprint-coding positions (rho = 0.53, Figure S7C), possibly due to the limited number of variants found. We also observed comparable rates of KZFP LoF variants in the NFE and AFR populations (rho = 0.67, Figure 5A), with 65 restricted to the NFE and 7 to the AFR subsets. While this difference might simply reflect a difference in sample sizes, some noteworthy outliers pointed to possible population-specific adaptation. We observed LoF variants in *ZNF114, ZNF426*, and *ZNF714* in the 63,369 NFE and in a group of 12,897 Finnish (FIN) individuals, whereas none were found in the 12,020 AFR (Figure 5B). According to GTEx, *ZNF714* and *ZNF426* are expressed in most tissues, whereas *ZNF114* expression is restricted to the thyroid (Figure 5C). The genomic targets of ZNF426 are yet to been identified. The results of ChIP-seq experiments performed in 293T cells overexpressing tagged forms of ZNF114 and ZNF714 [1] revealed rare peaks for the former, while the latter was found significantly enriched at a subset of HERVK integrants (Figure 5D). Gene ontology (GO) enrichment analyses yielded significant biological process terms only for ZNF114, pointing to its involvement in chromatin silencing and wound healing (Figure S8).

**Figure 5.**
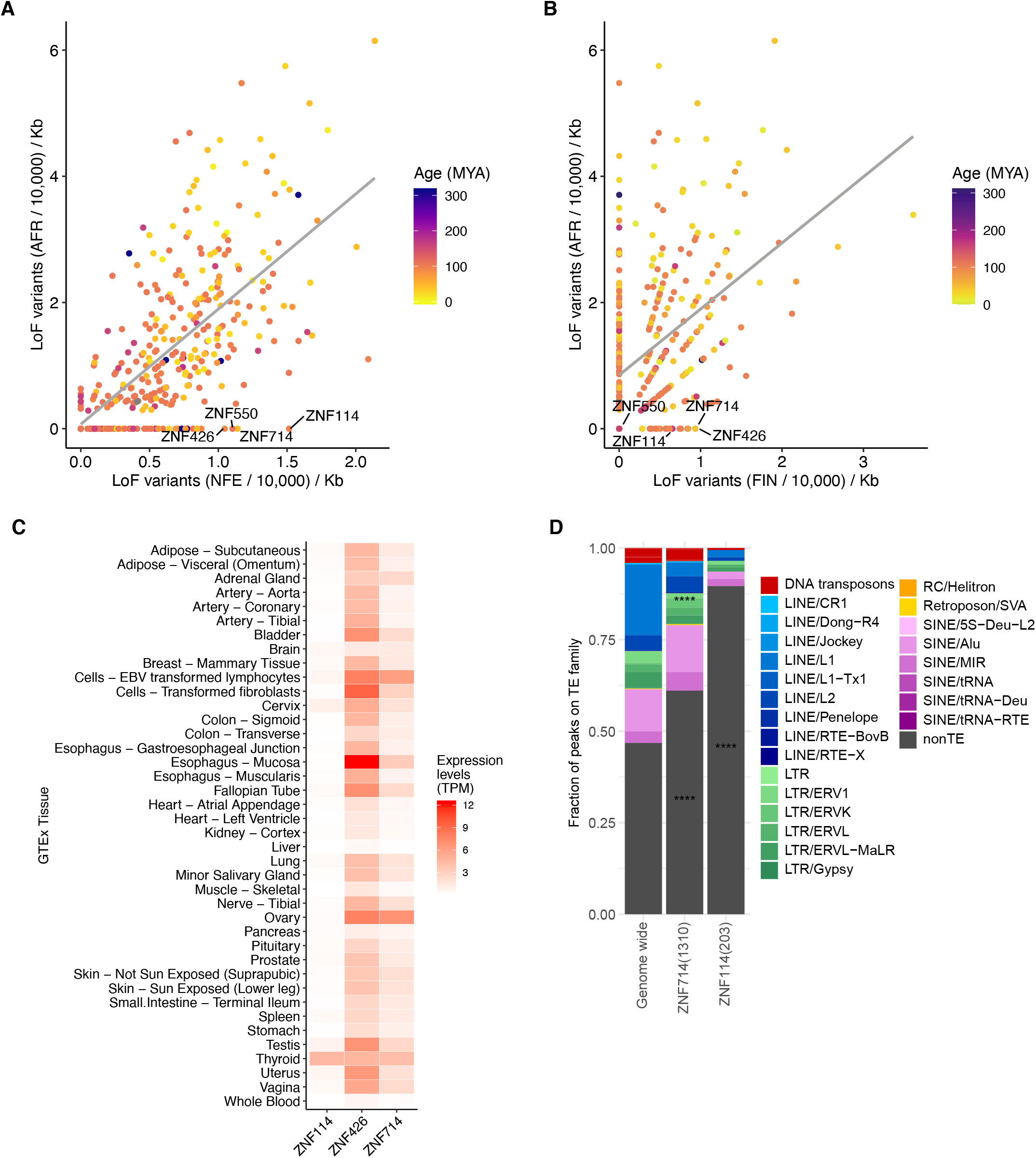
Population-specific constraint of human KZFPs. (A) Frequency of gnomAD-catalogued LoF variants in Africans (AFR) and Non-Finnish Europeans (NFE) per 10,000 individuals and kb of coding sequence. Grey line indicates the linear fit of the data. Common major KZFP outliers are annotated by name. (B) Same as in (A) crossing similarly sized groups of Finnish (FIN) and African individuals. (C) Expression levels in transcripts per million (TPM) for *ZNF114, ZNF426*, and *ZNF714* across 40 tissues. (D) Enrichment of the genomic TE targets of ZNF114 and ZNF714 from ChIP-exo peaks in 293T cells compared to the genome wide distribution of TEs. Stars indicate significant enrichments and grey bars non-TE genomic targets.

## Discussion

KZFPs are amongst the fastest evolving vertebrate genes. Amongst the 361 protein-coding family members with good sequencing coverage in the human genome included in this study, 87% were placental mammalian-restricted (≤ 105 MYO), and 33% were primate-specific, down from 45% for all putative primate-specific KZFP coding sequences as these comprise numerous annotated pseudogenes [1]. The present work demonstrates that the evolutionary conservation of individual KZFPs correlates with the genetic constraint imposed on their coding sequences. Older KZFPs display lower degrees of genetic variation in the human population than their more recent counterparts, in particular at positions encoding amino acids predicted to dictate the DNA binding specificity of their products. Recent studies indicate that highly conserved KZFPs, some of which can be traced back to the emergence of tetrapods, only rarely bind to recognizable TE-derived sequences. They often associate with proteins that evoke essential functions such as RNA metabolism or chromosome architecture, and are usually devoid of paralogs [1,4]. Our observation that these genes are under severe coding constraints is therefore not surprising. In contrast, younger KZFPs almost universally target TEs, often have paralogs with zinc fingerprints suggestive of at least partly overlapping sets of genomic targets, and display a KAP1-centered protein interactome primarily consistent with transcriptional repression [1,4]. The greater genetic variation observed in the human population at positions determining the genomic targets of these recently emerged KZFPs may thus be explained by functional redundancy between closely related family members or by the absence of TEs forcing fixation of at least part of their ZF-coding sequences. Conversely, this also suggests that genetic variations in KZFP gene sequences, together with differences in the genomic distribution and sequence of their TE targets, participate in the phenotypic diversification of modern humans.

Differentials in coding constraint greatly varied within identified sets of unequivocal KZFP paralogs, being very narrow in some cases (e.g., *ZNF75A* and *ZNF75D; ZFP30* and *ZFP82*) and quite broad in others (e.g., *ZNF160* and *ZNF665; ZNF181* and *ZNF302*). No single parameter could account for these differences. For instance, ZNF75A and ZNF75D both recognize the 3’end of KZFP genes, while ZFP30 and ZFP82 respectively bind LINEs and SINEs, that is, completely distinct sets of genomic targets. As well, *ZNF679-ZNF716* and *ZNF600-ZNF611* are two pairs of evolutionarily recent (<20 MYO) paralogs, yet they present with coding constraint differentials that are negligible for the former and pronounced for the latter. Still, it is noteworthy that for paralogs of detectably distinct ages, the older one was found to be generally more constrained than its duplication product, recapitulating a trend noted for the KZFP family as a whole. This supports a general model whereby KZFP paralogs diversify the trans-regulatory space available for coping with new genomic targets resulting from genetic drift in the TE load of the host organism, while the corresponding parent genes keep controlling pre-existing cis-acting regulatory sequences, ensuring that physiology is maintained. This mechanism also explains the high turnover in the pool of a lineage’s KZFP genes, even if only a fraction of newcomer genes is positively selected.

Our data indicate that evolutionary forces have also shaped the KZFP gene pool of modern humans in the recent past. It is generally considered that *Homo sapiens* started durably settling out of Africa some 60,000 years ago and that the genetic diversity of these early migrants was reduced compared to individuals left behind owing to their comparatively smaller number and due to selection bottlenecks. Accordingly, for most genes, the number of observed polymorphisms is expected to be either equal or greater in individuals of African descent compared with Europeans or Asians, a trend verified for the vast majority of human KZFP genes. Thus, it is noteworthy that at least three family members, *ZNF114*, *ZNF426*, and *ZNF714*, all three several tens of million years old, exhibit the opposite pattern with LoF mutants absent in individuals of African descent but observed in a European subpopulation of similar size. The first, *ZNF114*, has not been functionally examined previously, but is expressed mainly in the thyroid, with a GO enrichment analysis pointing to its involvement in wound healing. The second, *ZNF714*, has previously been associated with insulin resistance [39], a trait that exhibits substantial population-specific differences with Africans having significantly lower insulin sensitivity and higher insulin responses than Europeans [40]. The third, *ZNF426*, was previously found to bind to a Kaposi’s sarcoma-associated herpesvirus (KSHV) transactivator and to reduce as a consequence the transcriptional activity of this virus, qualifying ZNF426 as an innate antiviral restriction factor [41]. As KHSV is rare in Europe and the Americas but endemic in Africa [42], it is tempting to speculate that the presence of this pathogen contributed to the maintenance of functional *ZNF426* alleles in the African population, and that this selective pressure was alleviated for descendants of humans having long left this continent. Further exploration of these various hypotheses will require the analyses of greater numbers of genomes from individuals of non-European ancestry to rule out genetic drift as the source of these population variation differences, as well as functional experiments directly addressing the roles of ZNF114, ZNF426, ZNF714 and possibly other KZFPs displaying differential coding constraints in various human subgroups.

## Conclusions

Overall, we provide evidence of the genome-wide selection and constraint of human KZFPs, even within functionally important protein sites, and their role in shaping the current genetic variation within this large family of genome-regulating proteins with implications for the biology of modern humans.

## List of abbreviations

AFR: African/African-Americans
C2H2: Cys2-His2
FIN: Finnish
GO: Gene ontology
KAP1: KRAB-associated protein 1, also known as TRIM28
KRAB: Krüppel-associated box
KSHV: Kaposi’s sarcoma-associated herpesvirus
KZFP: Krüppel-associated box (KRAB) zinc finger protein (KZFP)
LoF: Loss-of-function
MYA: Million years ago
MYO: Million years old
NFE: Non-Finnish Europeans
TE: Transposable element
TEeRS: transposable element-embedded regulatory sequences
ZF: Zinc finger

## Declarations

### Availability of data and materials

The datasets supporting the conclusions of this article, the KZFP genomic map and calculated genetic constraint scores are available in the Figshare repository with the accession https://dx.doi.org/10.6084/m9.figshare.11663475 and https://dx.doi.org/10.6084/m9.figshare.11733333, respectively.

### Authors’ contributions

C.W.T., J.F., P.T., and D.T. contributed to the conception and design of the study. C.W.T., E.P., J.D.T., A.C., J.F., P.T., and D.T. contributed to the analysis and interpretation of data. C.W.T., J.D.T. and D.T. contributed to the drafting the manuscript. All authors reviewed and approved the final manuscript.

## Acknowledgements

Not applicable

