## Supplementary figures for "Ongoing evolution of KRAB zinc finger protein-coding genes in modern humans"

### Supplementary Material

#### Figure S1

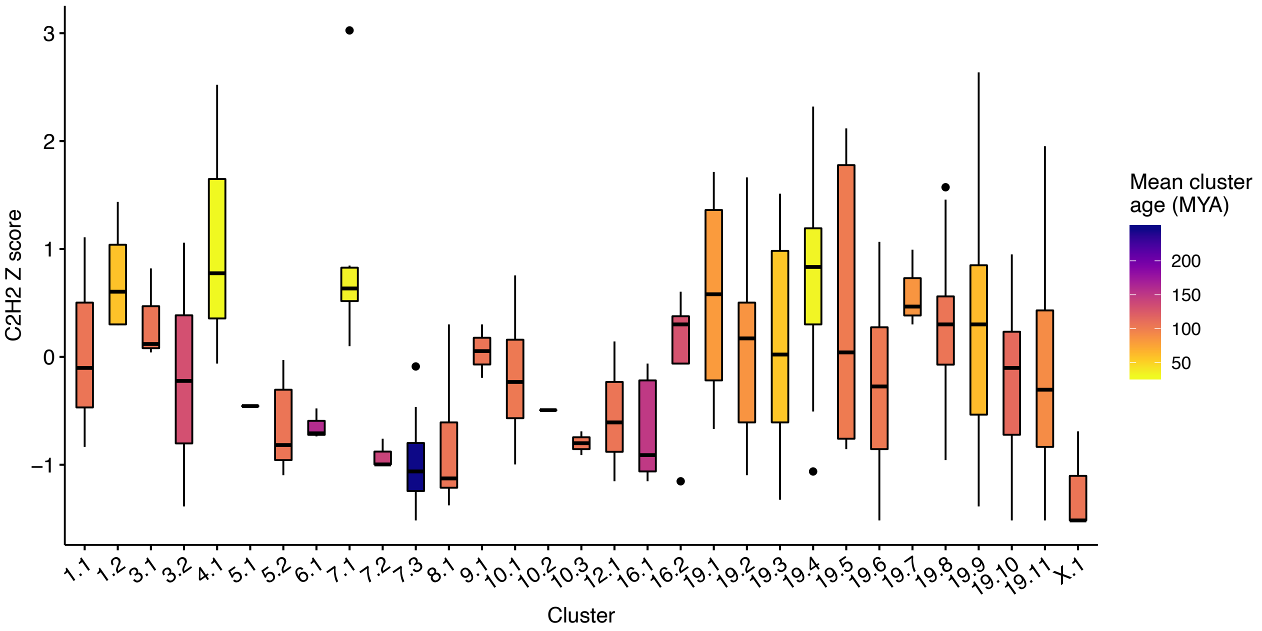


**Figure S1. Variable constraint within each KZFP cluster.** The variable levels of constraint in the cysteine and histidine levels across KZFP clusters, colored by the mean age (MYA) of the KZFPs within each genomic cluster.

#### Figure S2

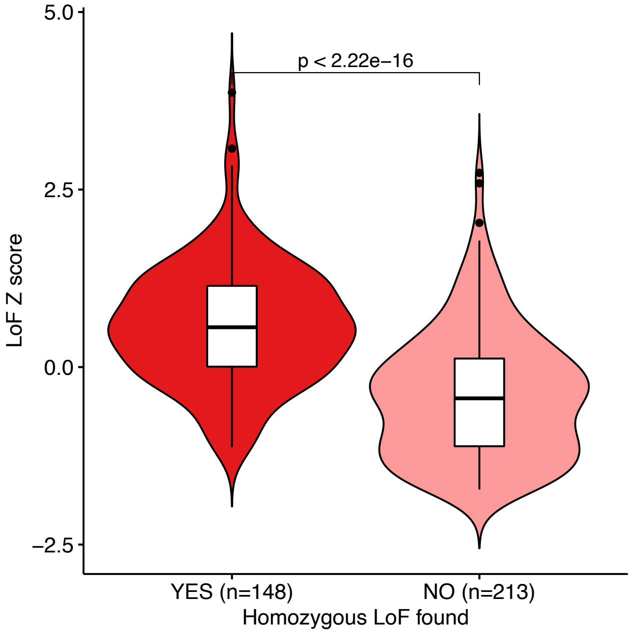


**Figure S2. Relationship between LoF constraint and presence of homozygous LoF variants.** KZFPs in which homozygous LoF variants were found are less constrained than KZFPs were only heterozygous LoF variants were identified. The p-value were obtained with the Wilcoxon rank sum test.

### Figure S3

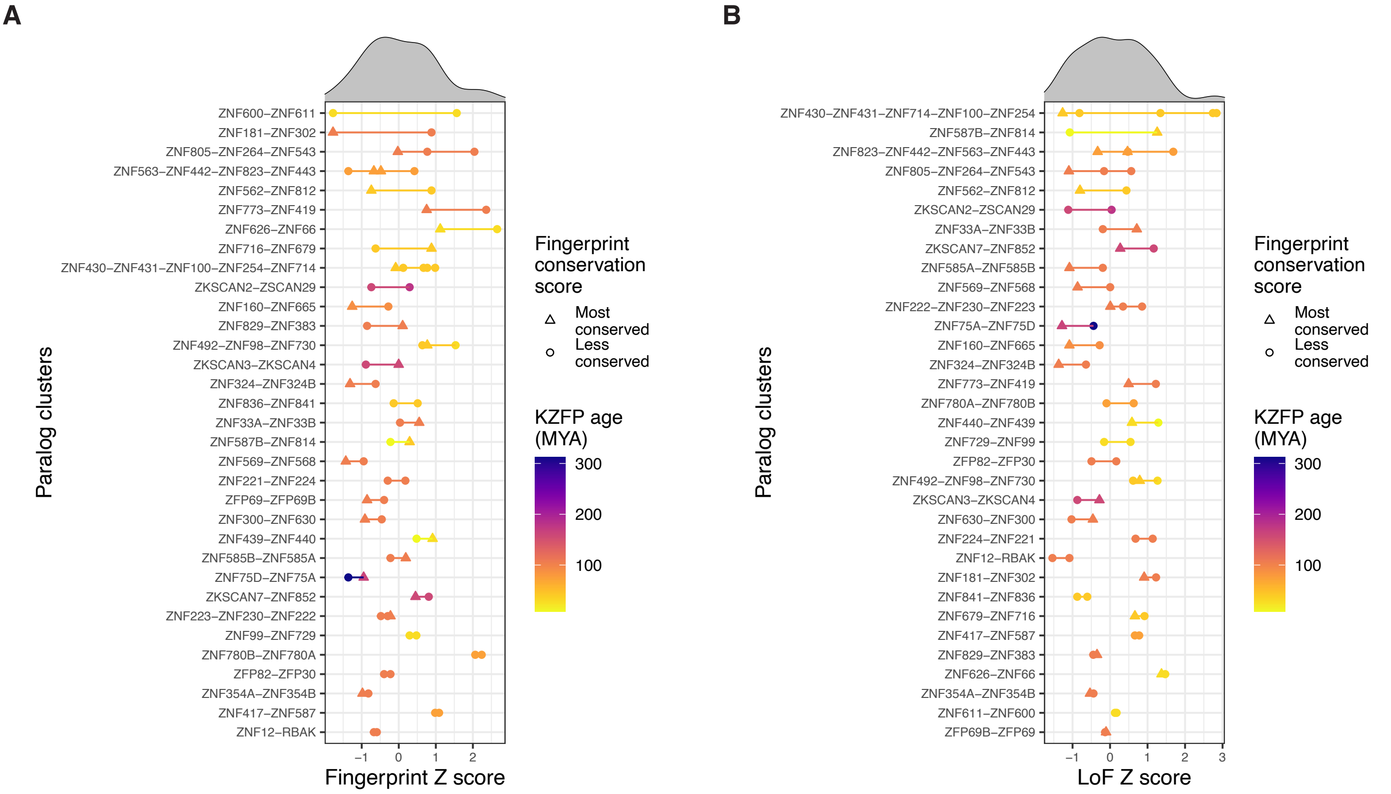


**Figure S3. Fingerprint and LoF constraint within paralog pairs.** (A) Fingerprint constraint within paralog pairs, sorted with the pairs exhibiting the largest differences on top. Colors represents the estimated age of the KZFPs. The KZFP within each paralog pair with the most conserved fingerprint across evolutionary time and species is marked by a triangle. (B) LoF constraint within paralog pairs.

### Figure S4

**A**

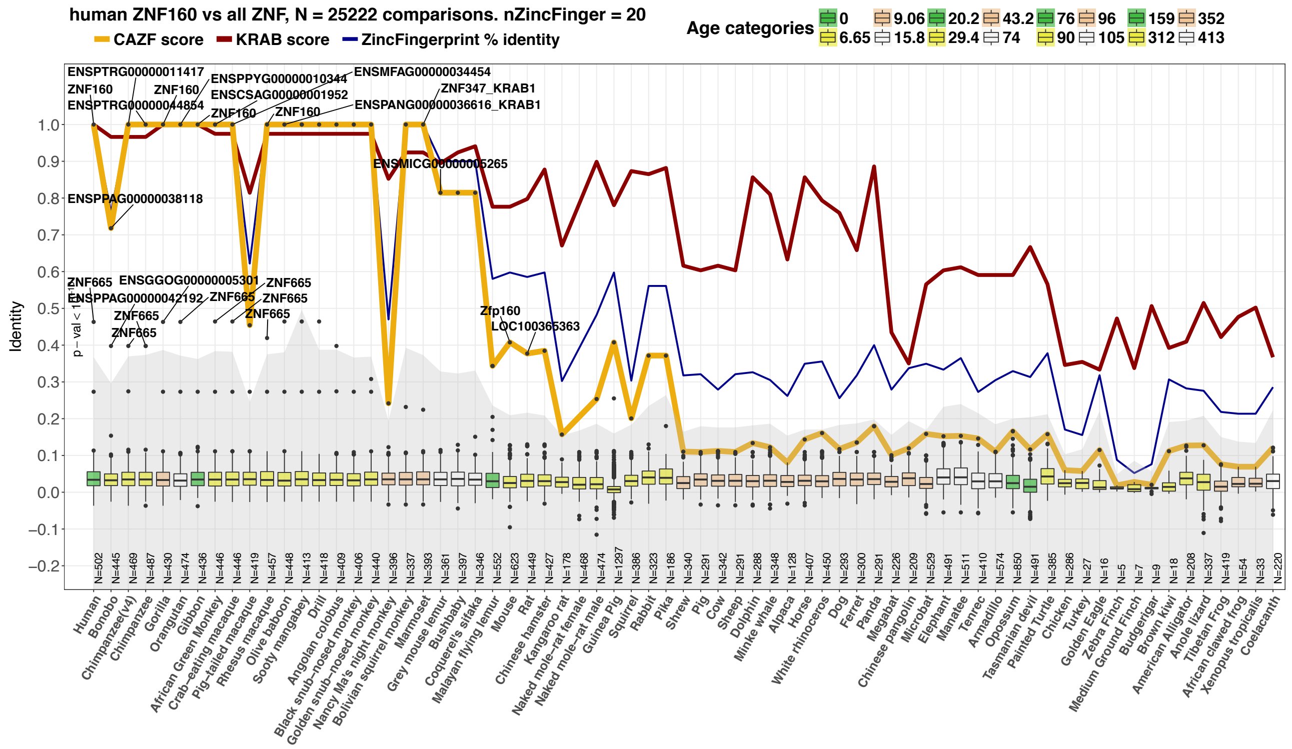


**B**

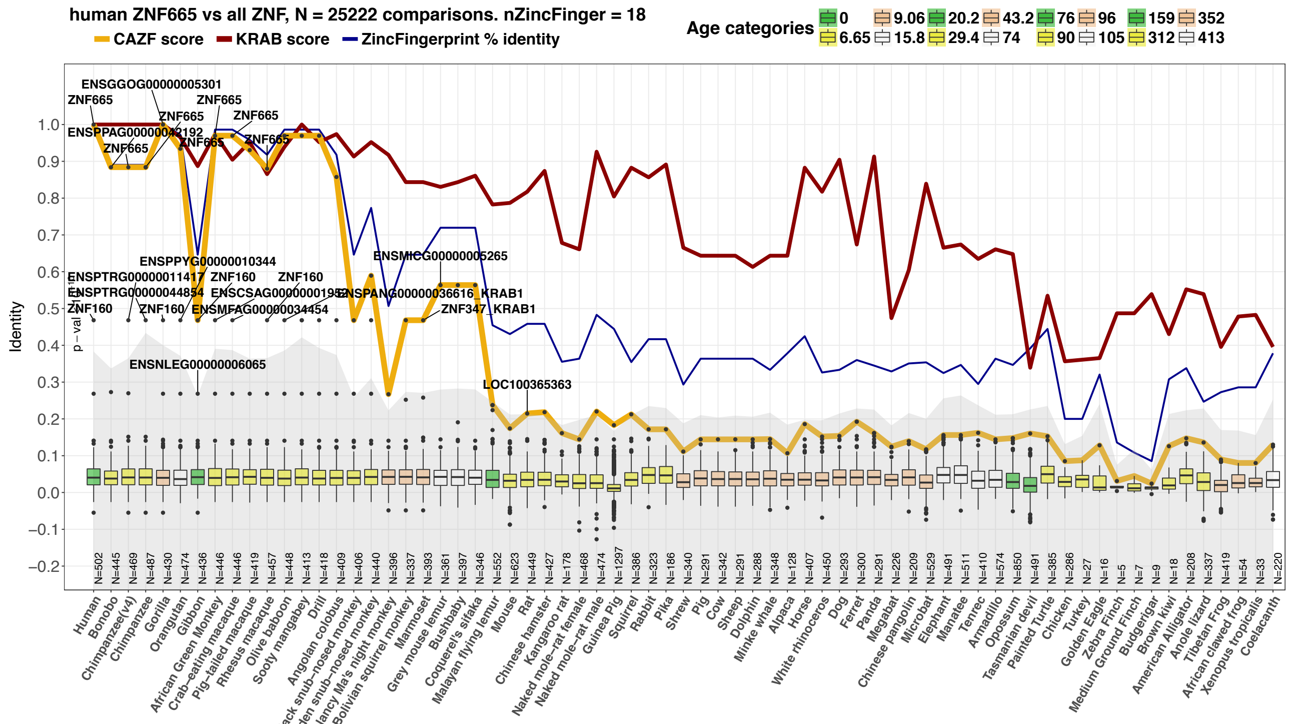


**Figure S4. Zinc fingerprint conservation across species for ZNF160 and ZNF665.** (A) Visualization of the similarity (percentage) between the modern human zinc fingerprint of ZNF160 (blue line) and that across evolutionary time and species. Age categories are based on the estimated time of divergence. The CAZF score (yellow line) indicates the zinc fingerprint evolutionary conservation score. Red line indicates the similarity of the KRAB domains. (B) ZNF665 fingerprint similarity across species.

#### Figure S5

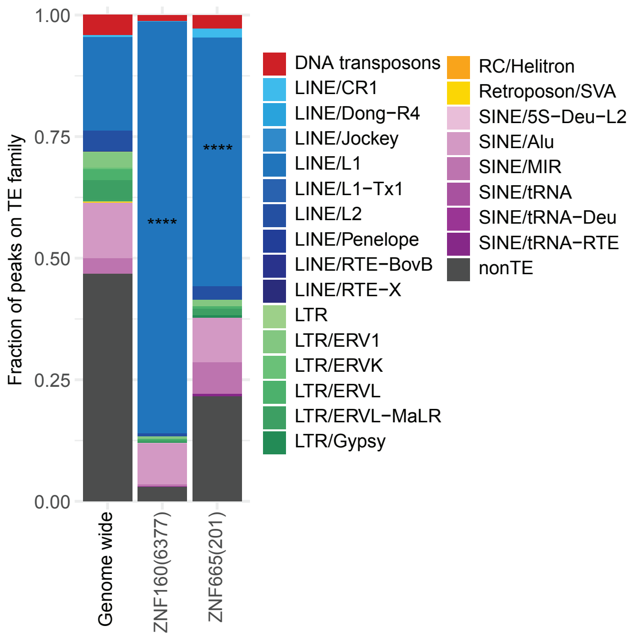


**Figure S5. Similar genomic targets for ZNF160 and ZNF665.** TE target enrichment analyses from ChIP-exo of ZNF160 and ZNF665 relative to the genome wide distribution of TEs. Stars indicates statistically significant enrichments. The numbers in brackets after the KZFP name indicates the number of peaks found for each ChIP-exo. The majority of the peaks for both ZNF160 and ZNF665 falls within LINE1 elements.

#### Figure S6

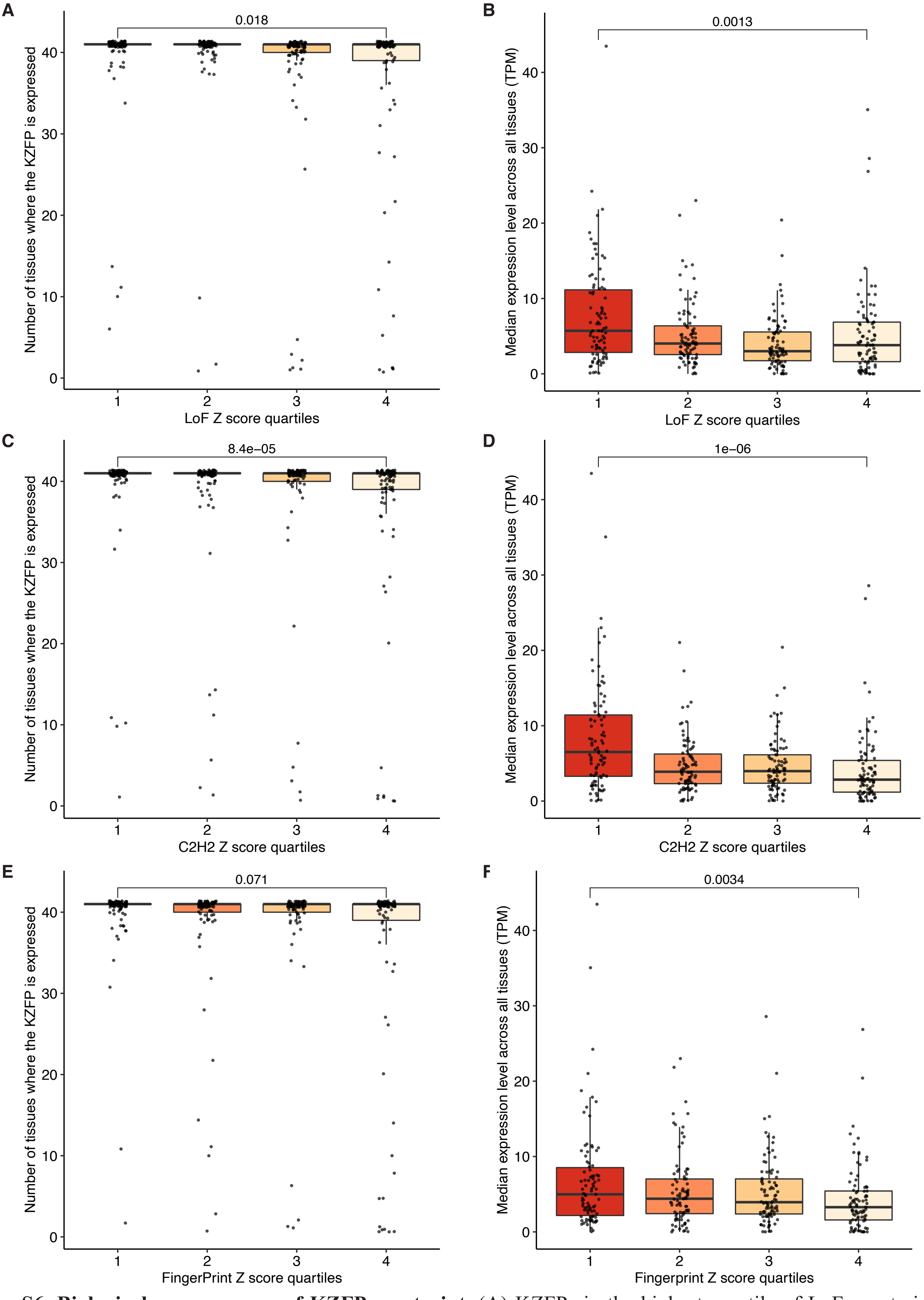


**Figure S6. Biological consequences of KZFP constraint**. (A) KZFPs in the highest quartile of LoF constraint are expressed in more tissues than KZFPs in the lowest quartile. (B) The mean expression levels of KZFPs in the first quartile of LoF constraint is higher than for those KZFPs in the fourth constraint quartile (lowest constraint). (C) The most C2H2 constrained KZFPs are also expressed in more tissues and (D) at higher levels. (E) Constraint of the fingerprint residues is also associated with more ubiquitous expression (F) and higher levels of expression. All p-values are from Wilcoxon rank sum tests.

### Figure S7

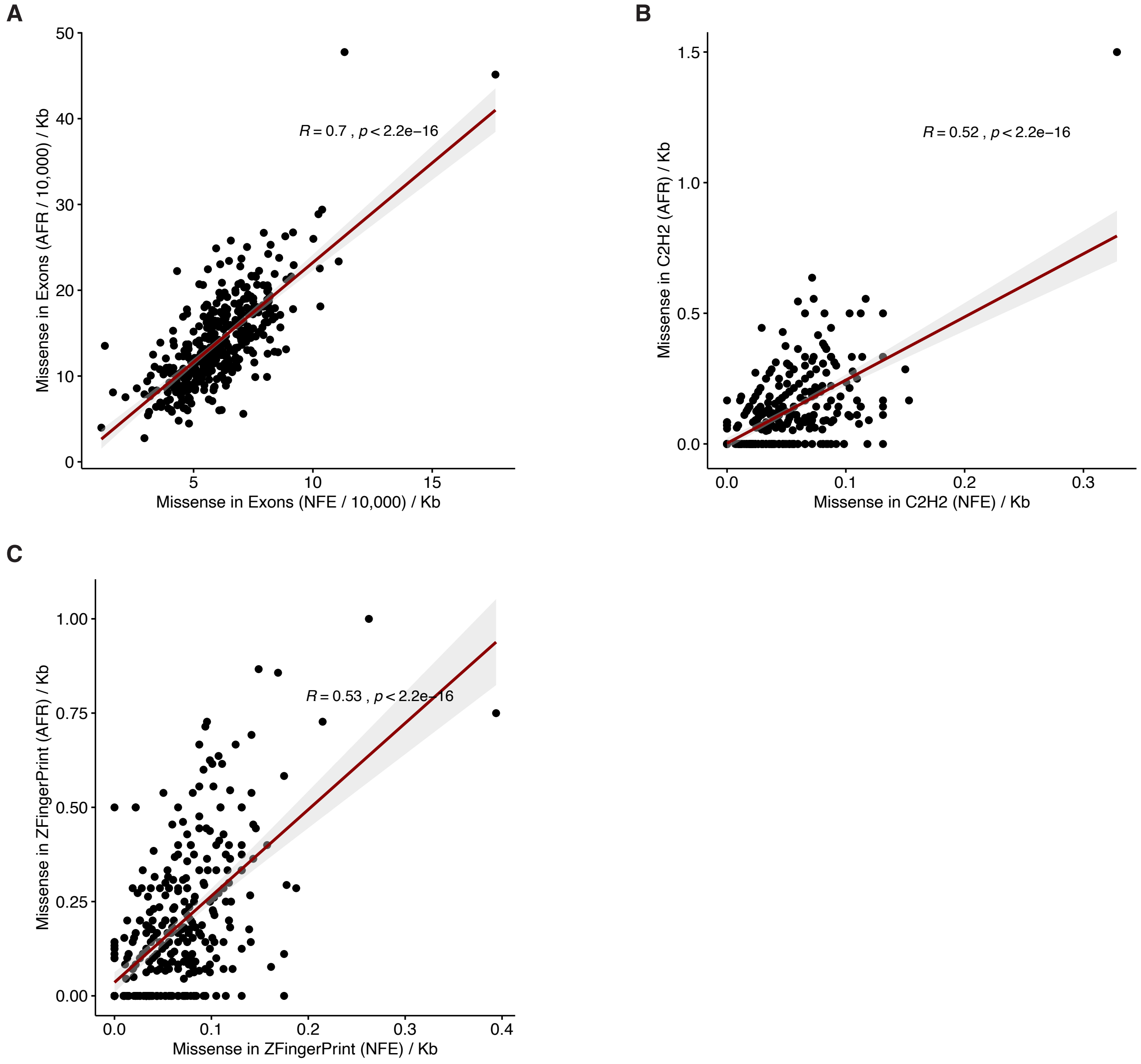


**Figure S7. Correlations across population specific constraints.** (A) Spearman correlations between number of observed missense variants per 10,000 individuals in non-Finnish Europeans (NFE) and Africans (AFR) within KZFPs, normalized by the length of the coding sequence. (B) Correlation between observed missense variants in the C2H2 positions within each KZFP between NFE and AFR individuals. (C) Correlation between observed missense variants in the zinc fingerprint of each KZFP between NFEs and AFRs.

### Figure S8

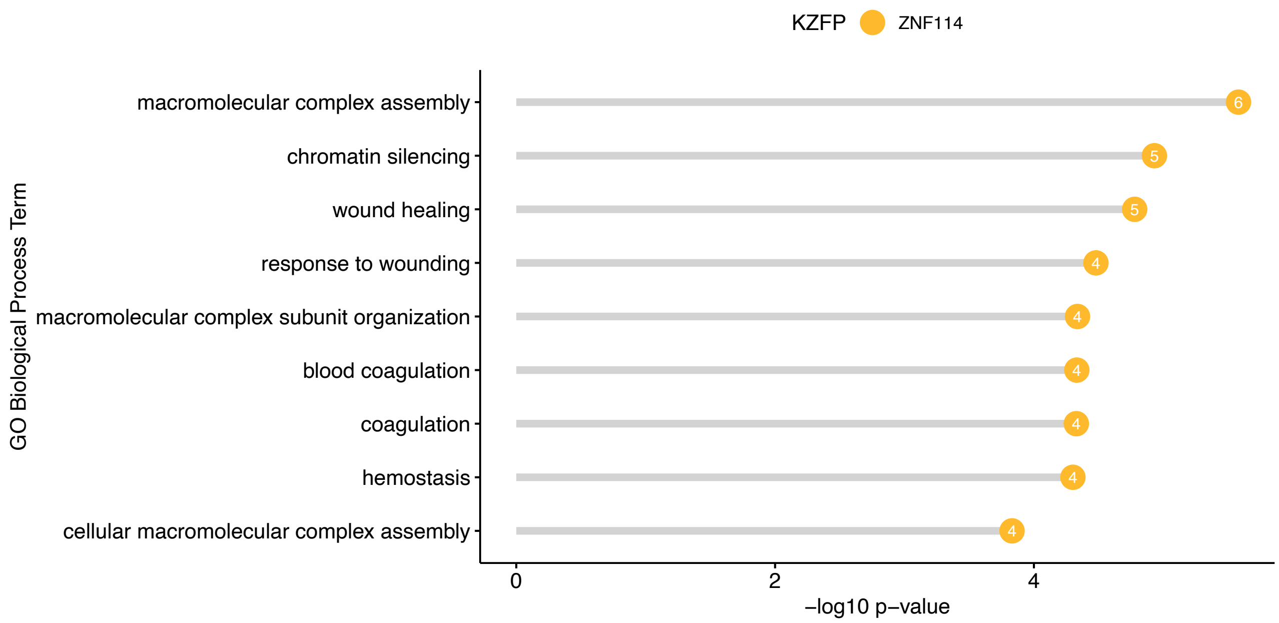


**Figure S8. Enrichment of GO Biological Process Terms for ZNF114.** Significant GO enrichment terms, as determined by GREAT, for ZNF114 shown with their raw -log10 binomial p-values.
